# Coexistence of complete competitors with fitness inequality

**DOI:** 10.1101/160986

**Authors:** Lev V. Kalmykov, Vyacheslav L. Kalmykov

**Affiliations:** Institute of Theoretical and Experimental Biophysics, Russian Academy of Sciences, Pushchino, Moscow Region 142290, Russian Federation; Pushchino State Institute of Natural Sciences, Pushchino, Moscow Region 142290, Russian Federation; Institute of Cell Biophysics of Russian Academy of Sciences, Pushchino, Moscow Region 142290, Russian Federation

## Abstract

There was a long standing contradiction between formulations of the competitive exclusion principle and natural species richness, which is known as the biodiversity paradox. Here we investigate a role of fitness differences in coexistence of two completely competing species using individual-based cellular automata. According to the classical formulations of the competitive exclusion principle such coexistence is impossible. Earlier we found that coexistence of complete competitors is possible with a 100% difference in fitness, but only under certain initial conditions. Here we verify a hypothesis that completely competing species may coexist with less than 100% difference in fitness regardless of different initial location of competing individuals in the ecosystem. We have found a new fact that two aggressively propagating complete competitors can stably coexist in one limited, stable and homogeneous habitat, when one species has some advantage in fitness over the other and all other characteristics of the species are equal, in particular any trade-offs and cooperations are absent. This fact is established theoretically on the rigorous model. The found competitive coexistence occurred regardless of the initial location of individuals in the ecosystem. When colonization of free habitat started from a single individual of each species, then the complete competitors coexisted up to 31% of their difference in fitness. And when on initial stage half of the territory was probabilistically occupied, the complete competitors coexisted up to 22% of their difference in fitness. These results additionally support our reformulation of the competitive exclusion principle, which we consider as resolving of the biodiversity paradox.

## INTRODUCTION

Specific mechanisms of interspecific competition still remain insufficiently understood (Clark, 2009; Clark, 2012; Sommer, 1999; Tilman, 1987; Whitfield, 2002). This principle states that species competing for one and the same limiting resource in one homogeneous, limited and stable habitat cannot coexist. A brief formulation of the principle was: *“Complete competitors cannot coexist”*(Hardin, 1960). In theory, according to the competitive exclusion principle, complete competitors cannot coexist, but in practice, there are many examples of such coexistence: tropical rainforest, coral reefs, plankton communities (Sommer, 1999). This contradiction is known as the biodiversity paradox. Resolving the diversity paradox became the central issue in theoretical ecology (Lehman and Tilman, 1997). The urgent tasks of biodiversity conservation became additional motivation of the long-standing biodiversity debates. As biodiversity of trophically related species is the fact, then the problem of solving the paradox is reduced to verification of the competitive exclusion principle. This verification is necessary for elimination of the biodiversity paradox (Sommer, 1999). According to Hardin, the particular difficulty of the problem stems from the fact, that the principle cannot be proved or disproved by empirical facts, but only by theory (Hardin, 1960). Thus, verification of the principle became the great challenge for mathematical modelling.

The most well-known model of interspecific competition is the Lotka-Volterra model. This model is based on differential equations and that is why it is nonmechanistic or phenomenological (Huston et al., 1988; Kalmykov and Kalmykov, 2013; Kalmykov and Kalmykov, 2016; Tilman, 1987). This model is of black-box type because, despite the fact that it is deterministic, it does not model neither local interactions nor part-whole relationships. This model predicts stable coexistence of two similar species when, for both species, an interspecific competition is weaker than intraspecific one. This interpretation follows directly from the model. However, the further interpretation of this interpretation in form of the competitive exclusion principle has no rigorous justification under itself.

The unified neutral theory of biodiversity and biogeography (neutral theory or UNTB) was proposed as an attempt to solve the biodiversity paradox (Hubbell, 2001). The neutral theory proposed to replace the study mechanisms of interspecific interactions by statistical predictions of species presence-absence. This way is based on the assumptions which are clearly not true. Unreality of the hypothesis of ecological equivalence is obvious and for Hubbell with colleagues, as they assert that *“the real world is not neutral”* (Rosindell et al.). The long-standing debates on the biodiversity paradox as “competitive exclusion principle versus natural biodiversity” has been substituted for theoretical debates *“neutrality versus the niche”* (Kalmykov and Kalmykov, 2016; Whitfield, 2002). Understanding of biodiversity mechanisms should be based on mechanistic models. The UNTB is not based on a mechanistic model, – *“it is just a statement of ignorance about which species can succeed and why”* (Clark, 2009; Clark, 2012). Tilman considered that *“experiments that concentrate on the phenomenon of interspecific interactions, but ignore the underlying mechanisms, are difficult to interpret and thus are of limited usefulness”* (Tilman, 1987). Thus, in order to solve this problem it is necessary to create a mechanistic model of species coexistence.

There are individual-based cellular automata models of interspecific competition (Reichenbach et al., 2007; Silvertown et al., 1992), but they are used rather for fitting to phenomenology of experimental data than for understanding of fundemental ecological laws. In our models we have used a physical semantics of ecosystem dynamics (Kalmykov and Kalmykov, 2013, 2015a, b). We transform this semantics to a set of cellular automata rules. An elementary object of our models is a microecosystem. This is the basic abstrast autonomous object. Autonomous dynamics of this object is characterized by a set of states and the diagram of their sequential transitions between states in accordance with physical extremal principles. Following Tansley, we consider individuals with their *‘‘special environment, with which they form one physical system’’* (Tansley, 1935). A microecosystem includes a single microhabitat and is able to provide a single accommodation of one individual, but it cannot provide its propagation. After an individual’s death, its microhabitat goes into a regeneration (refractory) state, which is not suitable for immediate occupation. In our models vegetative propagation is carried out due to resources of an individual’s microhabitat by placing its vegetative primordium of a descendant in a free microhabitat of an individual’s neighbourhood. A plant ecosystem may be considered *“as a working mechanism”* which *“maintains and regenerates itself”* (Watt, 1947). Our model demonstrates such the mechanism. From a more general physical point of view we model an active (excitable) media with population autowaves (travelling waves, self-sustaining waves) (Kalmykov and Kalmykov, 2013; Krinsky, 1984; Zaikin and Zhabotinsky, 1970). An active medium is a medium that contains distributed resources for maintenance of autowave propagation. An autowave is a self-organizing and self-sustaining dissipative structure. An active medium may be capable to regenerate its properties after local dissipation of resources. In our model, propagation of individuals occurs in the form of population autowaves. We use the axiomatic formalism of Wiener & Rosenblueth (Wiener and Rosenblueth, 1946) for modelling of excitation propagation in active media, where. In accordance with this formalism rest, excitation and refractoriness are the three successive states, where the rest state corresponds to the free state of a microhabitat, the excitation state corresponds to the life activity of an individual in a microhabitat and the refractory state corresponds to the regeneration state of a microhabitat.

Taking into account the refractory state of microhabitat expands possibilities of individual-based modelling of ecosystem dynamics because most ecological models still do not take into account regeneration of resources in plant communities. This problem was first discussed in general terms by Watt in 1947 (Grubb, 1977; Watt, 1947).

Reasoning from the first principles we model population waves as autowaves in active media. The opposite approache is based on reasoning by analogy and we consider it as phenomenological. The main difficulty of reasoning from first principles is the preliminary need to create a general theory of the modelled object (Kalmykov and Kalmykov, 2015a). The general theoretical representations allow formulating semantically exact rules of cellular automata.

Further models logically derive more and more details of objects under study as automatic logical inference from the first principles of ecosystem theory. They reflect state changes of each microecosystem. A feature of this model approach is its multilevel character. It simultaneously and in details models events at the micro-, mini- (meso-) and macro-levels of the system under study. The multilevel character of complex systems in nature is perhaps the greatest problem of their mathematical modelling. The implemented here approach overcomes this difficulty. The logical basis of our models may create an impression of their excessive simplicity and even of a toy character. Unlike such games as John Conway’s Game of Life, the rules of our models are based not on an arbitrary combination of parameters, but on the fundamental theoretical propositions (on the first principles). The multilevel organization of complex systems does not necessarily involve the use of a complex mathematical apparatus for their modelling. It is necessary to remember the Occam’s razor. If phenomena under study can be investigated on relatively simple models, why do we need to use a more heavy mathematics?

A large number of studies have been devoted to finding mechanisms that prevent the implementation of the competitive exclusion principle. There were found more than 120 (Palmer, 1994) of such mechanisms (Bennett and Bever, 2009; Dollhopf et al., 2003; Hastings, 1980; Hutchinson, 1961; Nowak, 2006; Wellborn, 2002): trade-offs; cooperative interactions between the competing species; genetic heterogeneity of populations; sexual reproduction, which increases genetic heterogeneity of populations; excess resources of ecosystems; heterogeneity of habitat; immigration, emigration, predation, parasitism, herbivory, diseases and other disturbances of populations; instability of the dominance of a species as a result of variability of environmental conditions; ontogenetic differences in fitness, in fecundity rates, in regeneration features of a habitat and in environmental requirements of competing species. However, identification of factors hindering the implementation of the principle of competitive exclusion had little effect on the formulation of the principle itself. At the same time, formulations of the principle gradually became more and more stringent:

- Survival of the fittest (Spencer, 1864);
- Complete competitors cannot coexist (Hardin, 1960);
- *n* species require at least *n* resources to ensure indefinite and stable equilibrium coexistence in a homogeneous environment (MacArthur and Levins, 1964; Narwani et al., 2009);
- No stable equilibrium can be attained in an ecological community in which some r of the components are limited by less than r limiting factors (Levin, 1970);
- Two populations (species) cannot long coexist if they compete for a vital resource limitation of which is the direct and only factor limiting both populations (Darlington, 1972).
- Given a suite of species, interspecific competition will result in the exclusion of all but one species. Conditions of the Principle: (1) Time has been sufficient to allow exclusion; (2) The environment is temporally constant; (3) The environment has no spatial variation; (4) Growth is limited by one resource; (5) Rarer species are not disproportionately favored in terms of survivorship, reproduction, or growth; (6) Species have the opportunity to compete; (7) There is no immigration (Palmer, 1994).

To carry out a rigorous verification of the principle, we made an attempt to find a mechanism for competitive coexistence under the extremely strict conditions of interspecific competition. In our experiments we excluded a possibility of implementing numerous mechanisms that prevent implementation of the principle (Palmer 1994). We excluded such factors as: environmental fluctuations, any trade-offs and cooperative interactions, immigration, emigration, predation, herbivory, parasitism, infectious diseases, inhomogeneities of habitat, genetic inhomogeneities of individuals within the competing populations. In addition, the competitors were identical consumers and competed for one limiting resource. Thus the conditions of our experiments were extremely unfavorable for competitive coexistence. As a result of our investigation we found two deterministic individual-based mechanisms of competitive coexistence (Kalmykov and Kalmykov, 2013, 2015a). *The first coexistence mechanism* is based on free resource gaps which help to eliminate direct conflicts of interest between competing species, and as result colliding population waves of different competing species interpenetrate through each other like soliton waves in physical systems (Kalmykov and Kalmykov, 2013). A possible mechanism of appearance of such gaps is moderate reproduction which was modeled through a hexagonal rosette-like cellular automata neighbourhood. *The second coexistence mechanism* is based on timely recovery of the limiting resource, its spatio-temporal allocation between competitors and limitations of habitat size (Kalmykov and Kalmykov, 2015a). This mechanism allows complete competitors coexist in spite of using standard hexagonal cellular automata neighbourhood which models aggressive propagation without gaps in population waves. However, this mechanism of indefinite coexistence was limited by the habitat size and initial location of individuals on the lattice. In any case, the principle must always be right and if there are exceptions, then either the principle is not true, or its wording is not correct. The revealed mechanisms of competitive coexistence violate the listed formulations of the competitive exclusion principle and as result we have reformulated it as follows:

> *If each and every individual of a less fit species in any attempt to use any limiting resource always has a direct conflict of interest with an individual of a most fittest species and always loses, then, all other things being equal for all individuals of the competing species, these species cannot coexist indefinitely and the less fit species will be excluded from the habitat in the long run* (Kalmykov and Kalmykov, 2013).

These strict clarifications in the formulation of the principle demonstrate a low probability of the realization of this principle in nature because the need to comply with all formulated conditions makes the competitive exclusion rather a rare event. The obvious rarity of implementation the competitive exclusion principle removes the paradox of biodiversity because it eliminates the contradiction between the principle and the observed natural facts. In addition, we generalized the reformulated principle, setting out conditions under which one competitor is able to displace all others:

> *If a competitor completely prevents any use of at least one necessary resource by all its competitors, and itself always has access to all necessary resources and the ability to use them, then, all other things being equal, all its competitors will be excluded* (Kalmykov and Kalmykov, 2015a; Kalmykov and Kalmykov, 2016).

This formulation allows us to interpret different mechanisms of coexistence in terms of availability of access to necessary resources. It helps us to understand a threat to biodiversity that may arise if one competitor will control use of all necessary resources of an ecosystem.

Nowadays, humankind is becoming such a global competitor for all living things (Cincotta et al., 2000; Ehrlich and Ehrlich, 2008; Vitousek et al., 1997). Overexploitation of limited necessary resources in result of unbridled human population growth causes a problem which is known as the ‘tragedy of the commons’ (Hardin, 1959, 1968).

Here, in order to investigate coexistence mechanisms in more depth, we study a hypothesis *that two completely competing species may coexist with smaller than 100% difference in fitness regardless of different initial location of individuals of competing species in the habitat*. Along with the model of colonization of free ecosystem starting from single individuals we investigate cases when the habitat is probabilistically populated by a lot of individuals at initial iteration.

Probabilistic populating of the habitat makes it possible to expand the investigated variants of the conditions of interspecies competition.

## METHODS

### Biological prototype of the model

A vegetative propagation of rhizomatous lawn grasses is the biological prototype of our model. One individual corresponds to one tiller. The tiller is a minimal semi-autonomous grass shoot that sprouts from the base. Rhizomes are horizontal creeping underground shoots using which plants vegetatively (asexually) propagate. Unlike a root, rhizomes have buds and scaly leaves. One tiller may have six rhizoms in the model. Six rhizoms per tiller correspond to aggressive vegetative propagating. A tiller with roots and leaves develops from a bud on the end of the rhizome. A populated microhabitat goes into the regeneration state after an individual’s death. The regeneration state of a site corresponds to the regeneration of microhabitat’s resources including recycling of a dead individual. All individuals are identical. Propagation of one individual offsprings leads to colonization of the uniform, homogeneous and limited habitat.

### Basics of the cellular-automata models

Let us define terms “microhabitat”, “minihabitat” and “macrohabitat” in more detales:

> *Microhabitat* is the intrinsic physical environment where a particular individual inhabits. A microhabitat is a totality of all environmental conditions which are necessary for an individual’s life (e.g. of a tiller) and place where regeneration of the resources is possible. A site represents a place which can be occupied by an individual autonomous agent (e.g. by an individual of a species) and contains necessary resources for its individual life. Microhabitat with an individual forms a microecosystem.
>
> *Minihabitat* is the intrinsic physical environment (specific natural home) where a particular individual of a species is able to propagate during one generation. Minihabitat with individuals forms a mini-ecosystem.
>
> *Macrohabitat* is the total environment where individuals may inhabit and propagate actually and potentially. Here, in the cellular automata model, the macrohabitat is the entire field of the cellular automata without individuals. Macrohabitat contains a complete set of all cells of the cellular automaton lattice. Macrohabitat with individuals forms an ecosystem.

In order to test our *hypothesis*, we use our individual-based cellular automata model of interspecific competition with the following ecological conditions (Figs. 1, 2):

1. A habitat has a limited size, is homogeneous (microhabitats are identical) and stable (i.e. environmental conditions are constant, – there are no any climate or abiotic resources fluctuations and, as a consequence, fitnesses of competitors remain unchanged;
2. Immigration, emigration, predation, herbivory, parasitism, infectious diseases and other disturbances are absent;
3. Competing species are per capita identical and constant in ontogeny, in fecundity rates, in regeneration features of habitats and environmental requirements.
4. Reproduction of the competing species occurs only vegetatively and the species are genetically homogeneous and stable (from generation to generation there is no change of heredity);
5. We model trophically identical consumers which may differ only in fitness;
6. There are no trade-offs and cooperative interactions between the competing species;
7. Offsprings of an individual may vegetatively occupy all nearest microhabitats. Such propagation is potentially aggressive and modeled by the hexagonal neighbourhood.

**Figure 1.**
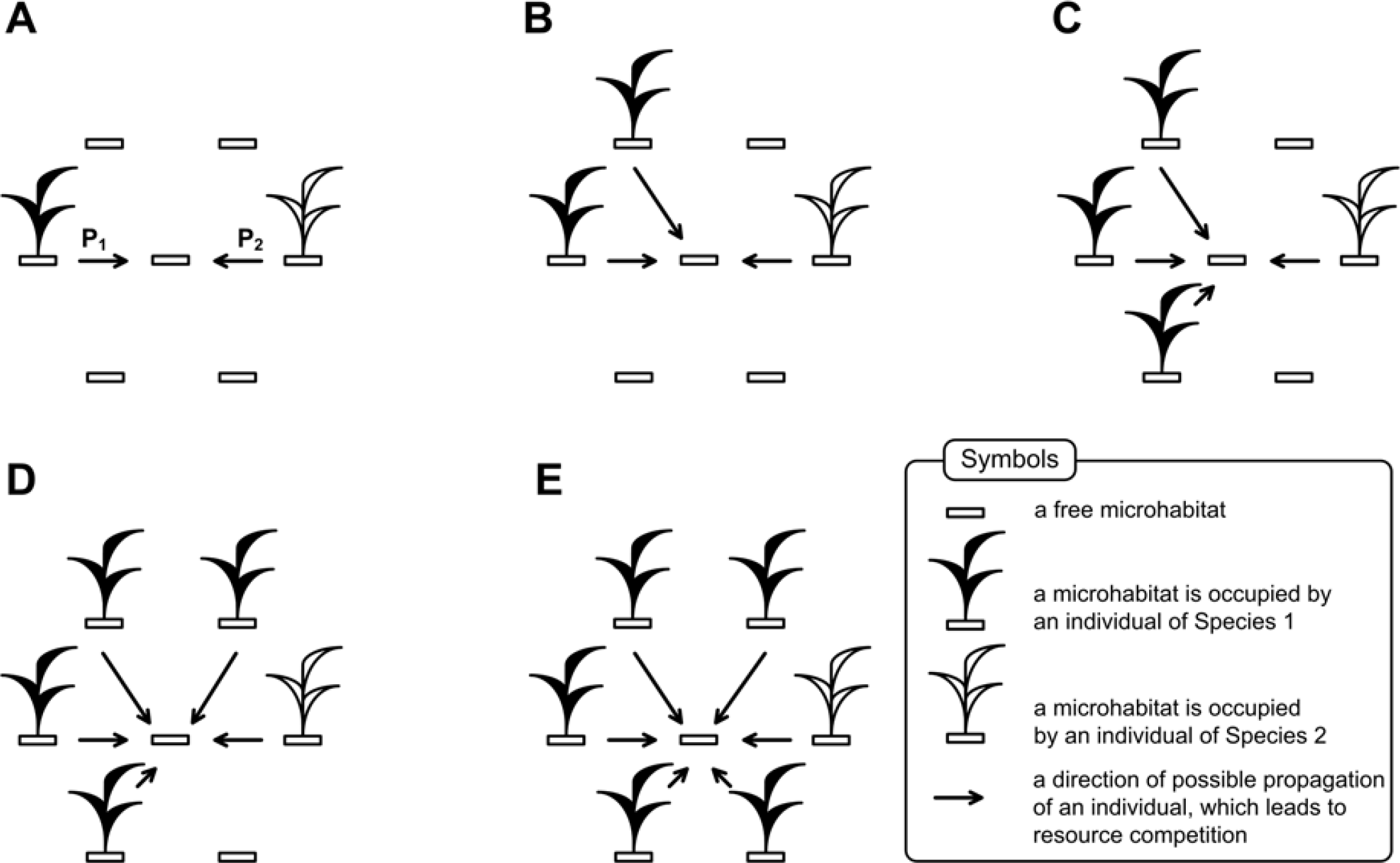
A schematic representation of interspecific competition for a free microhabitat. An increase in the number of individuals of a particular species that can occupy a free microhabitat does not increase chances of this species to occupy it. Parameters P_1_ and P_2_ represent probabilities of occupation of the free microhabitat by an offspring of Species 1 and Species 2, respectively. The maximum number of individuals, competing for the single microhabitat equals six (E). Chances to win in a direct conflict of interest are independent from number of competing individuals – parameters P_1_ and P_2_ are constant for all cases A-E.

**Figure 2.**
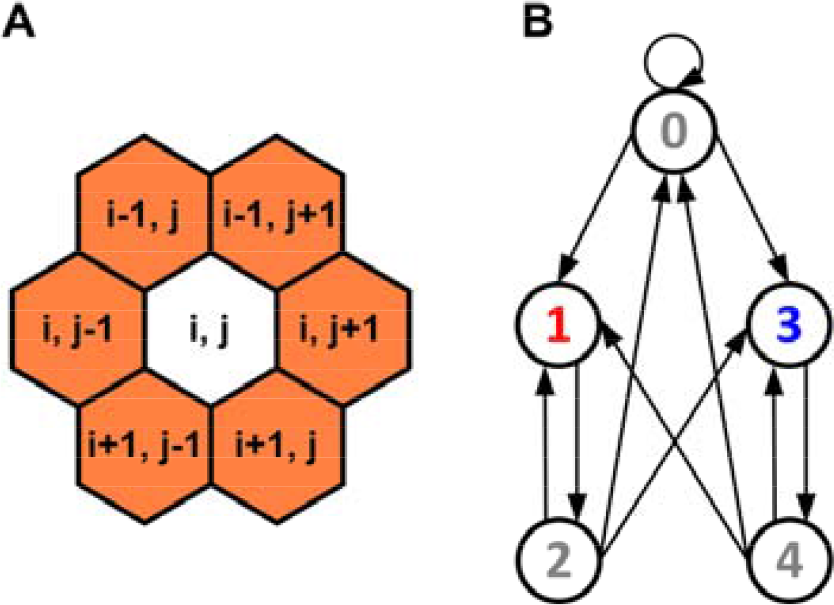
A description of the individual-based cellular automata model of an ecosystem with resource competition between two species. (A) Hexagonal neighbourhood. A central site of the neighbourhood is defined by the array element with index (i, j), where i and j are integer numbers. Neighbouring sites of the central site are defined by the array elements with indexes (i - 1, j), (i - 1, j + 1), (i, j + 1), (i + 1, j), (i + 1, j - 1), (i, j - 1). This neighbourhood allows to model vegetative propagation of individuals which theoretically may occupy all nearest sites, i.e. this is aggressive propagation. Possible local interactions between competing individuals are defined by this neighbourhood. (B) A directed graph of logical probabilistic transitions between the states of a lattice site. Each state and transition between states have a specific interpretation and represents a basic ecosystem ontology.

Fitness is a relative ability of the species to compete for environmental resources for propagation. In our models we define *individually oriented fitness – as a probability of reproductive success of an individual of the species in a direct competition for a free microhabitat*. In our model an individual of the Species 1 has a probability P_1_ to win in a direct conflict for resources, and an individual of the Species 2 has a probability P_2_ to win in the same conflict (Figs. 1, 2B). In all cases here, a microhabitat under the conflict of interest will always be occupied by an individual of one of the species, so P1 + P2 = 1. This excludes cases when a microhabitat may remain free at the next iteration in result of competition.

By changing fitness parameters P_1_ and P_2_ we look for a maximum value of fitness difference when competing species can coexist. The fitness difference is determined by the parameter P = P_1_ - P_2._ Starting from the deterministic case of competitive exclusion when P_1_ = 1, P_2_ = 0 and P = 1 we investigated most revealing cases of competitive coexistence with an increment of fitness difference which equals 0.01.

### The model of interspecific competition

Here we extend our logical cellular automata model of ecosystem with two competing species (Kalmykov & Kalmykov 2013; Kalmykov & Kalmykov 2015). We introduce probabilistic rules of competition to investigate the influence of fitness differences on the species coexistence. The model is based on the formalism of excitable medium and the concept of an individual’s intrinsic microecosystem (Kalmykov & Kalmykov 2015). A system of logical rules of transitions between the states of a lattice site of the cellular automata was formulated on the basis of a general theoretical concept of ecosystem. The entire cellular automaton simulates a whole ecosystem which autonomously maintains and regenerates itself. A two-dimensional hexagonal lattice is closed to a torus by periodic boundary conditions in order to avoid boundary effects. We use the hexagonal lattice because it most naturally implements the principle of densest packing of microhabitats. The hexagonal neighbourhood allows to model a potentially aggressive vegetative propagation of plants when offsprings of an individual may occupy all nearest microhabitats (Fig. 2A). Each site of the lattice simulates a microhabitat. A microhabitat contains resources for existence of a one individual of any species. A microecosystem is a microhabitat with an individual living in it. A microhabitat is the intrinsic part of environmental resources of one individual and it contains all necessary resources for its autonomous life. An individual can occupy a one microhabitat only. A life cycle of an individual lasts a single iteration of the automaton. All states of all sites have the same duration. Every individual of all species consumes identical quantity of identical resources by identical way, i.e. they are identical per capita consumers. Such species are complete competitors. Individuals are immobile in lattice sites and populations waves propagate due to reproduction of individuals (Movies S1-S3). The closest biological analogue is vegetative reproduction of rhizomatous lawn grass (Fig. 1).

A neighbourhood consists of a site and its intrinsically defined neighbour sites (Fig. 2A). All sites have the same rules for updating. A neighbourhood simulates an individual’s mini-ecosystem and determines the number of possible offsprings (fecundity) of the individual. The entire cellular automaton simulates a whole ecosystem (macroecosystem). The lattice of the cellular automaton simulates a habitat.

Here is a description of the states of a lattice site (Fig. 2B). Each site may be in one of the four states:

0—a free microhabitat which can be occupied by a single individual of any species;

1—a microhabitat is occupied by a living individual of the Species 1;

2—a regeneration state of a microhabitat after the death of an individual of the Species 1;

3—a microhabitat is occupied by a living individual of the Species 2;

4—a regeneration state of a microhabitat after the death of an individual of the Species 2.

Rules of transitions between the states of a site of the two-species competition model Fig.2):

0 → 0, A microhabitat remains free if there is no one living individual in the neighbourhood;

0 → 1, If in the cellular automata neighbourhood are individuals of both competing species, then the probability of this transition is defined by the parameter P_1_. If in the cellular automata neighbourhood is at least one individual of the Species 1 and there is no one individual of the Species 2, then this transition is always implemented;

0 → 3, If in the cellular automata neighbourhood are individuals of both competing species, then the probability of this transition is defined by the parameter P_2_. If in the cellular automata neighbourhood is at least one individual of the Species 2 and there is no one individual of the Species 1, then this transition is always implemented;

1 → 2, After the death of an individual of the Species 1 its microhabitat goes into the regeneration state;

2 → 0, After the regeneration state a microhabitat will be free if there is no one living individual in the neighbourhood;

2 → 1, If in the cellular automata neighbourhood are individuals of both competing species, then the probability of this transition is defined by the parameter P_1_. If in the cellular automata neighbourhood is at least one individual of the Species 1 and there is no one individual of the Species 2, then this transition is always implemented;

2 → 3, If in the cellular automata neighbourhood are individuals of both competing species, then the probability of this transition is defined by the parameter P_2_. If in the cellular automata neighbourhood is at least one individual of the Species 2 and there is no one individual of the Species 1, then this transition is always implemented;

3 → 4, After the death of an individual of the Species 2 its microhabitat goes into the regeneration state;

4 → 0, After the regeneration state a microhabitat will be free if there is no one living individual in its neighbourhood;

4 → 1, If in the cellular automata neighbourhood are individuals of both competing species, then the probability of this transition is defined by the parameter P_1_. If in the cellular automata neighbourhood is at least one individual of the Species 1 and there is no one individual of the Species 2, then this transition is always implemented;

4 → 3, If in the cellular automata neighbourhood are individuals of both competing species, then the probability of this transition is defined by the parameter P_2_. If in the cellular automata neighbourhood is at least one individual of the Species 2 and there is no one individual of the Species 1, then this transition is always implemented.

These logical statements include probabilistic parameters P_1_ and P_2_ which allow to investigate a role of fitness differences in coexistence of two agressively propagating and competing species. Parameter P_1_ reflects fitness of the Species 1 and parameter P_2_ reflects fitness of the Species 2. These cellular automata rules are realized on the three levels of organization of the complex system. A micro-level is modeled by a lattice site (micro-ecosystem). A mini-level of local interactions of a micro-ecosystem is modeled by the cellular automata neighbourhood (mini-ecosystem). A macro-level is modeled by the entire lattice (macro-ecosystem).

Different initial conditions may lead to formation of different spatio-temporal patterns and, as a result, to different dynamics of the system. We have investigated two situations on initial iteration – (i) when colonization of free habitat starting from a single individual of each species and (ii) when 25% of the territory was probabilistically populated by individuals of the Species 1, 25% of the territory was probabilistically populated by individuals of the Species 2 and 50% of the territory remained free. Here we show examples of initial patterns when the lattice consists of 50x50 sites (Fig. 3).

**Figure 3.**
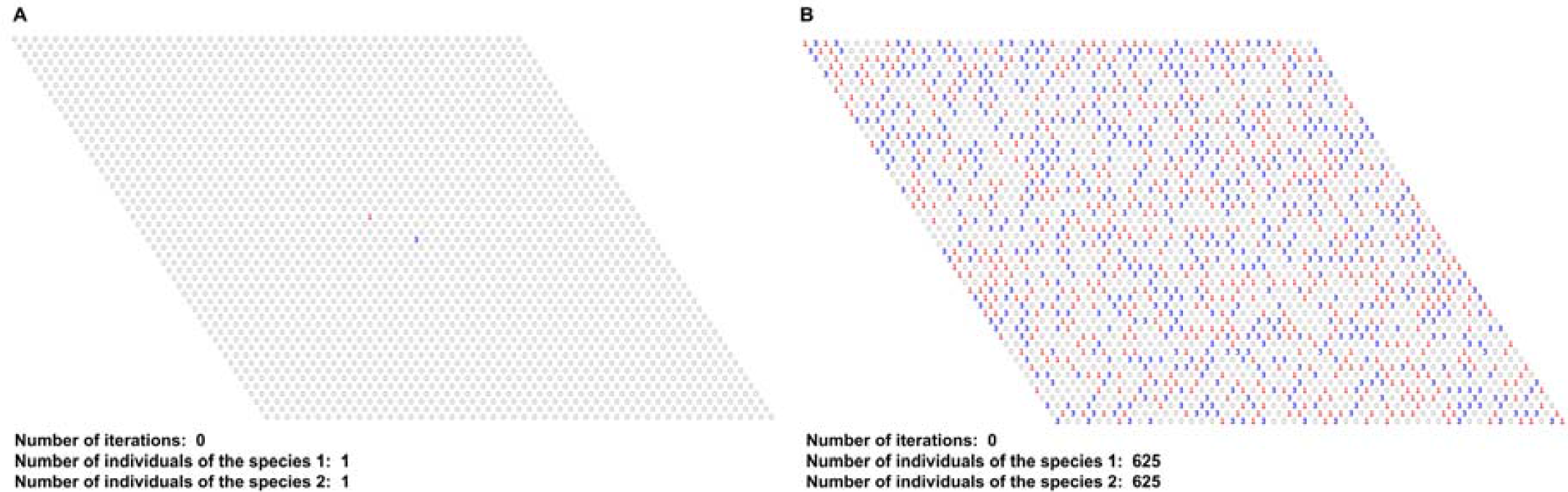
Examples of initial patterns. The two situations on initial iteration – (A) when colonization of free habitat starting from a single individual of each species and (B) when 25% of the territory was probabilistically populated by individuals of the Species 1, 25% of the territory was probabilistically populated by individuals of the Species 2. Cellular automata lattices consist of 50x50 sites and closed on the torus. (A) The cellular automata field was populated by single individuals of two competing species on initial iteration. (B) The field was probabilistically populated by 625 individuals of each of two competing species on initial iteration, i.e. 25% of the territory is occupied by individuals of the Species 1, 25% of the territory is occupied by individuals of the Species 2 and 50% of the territory is free.

## RESULTS AND DISCUSSION

Here we investigate the hypothesis that two completely competing species can coexist with smaller than 100% difference in fitness regardless of initial positioning of individuals in the habitat. We were changing fitness parameters P_1_ and P_2_ to find most specific cases of two species competition. We have identified most typical values of the parameters P_1_ and P_2_, which are responsible for coexistence and competitive exclusion (Fig. 4). We examined these most typical cases of population dynamics with one and the same initial positioning of individuals on the lattice. These are the case of competitive exclusion (Fig. 4A and Movie S1) and cases of competitive coexistence (Figs. 4B, 4C and Movies S2, S3). Figures 4A-4C demonstrate population dynamics in computer experiments presented in Movies S1-S3, respectively.

**Figure 4.**
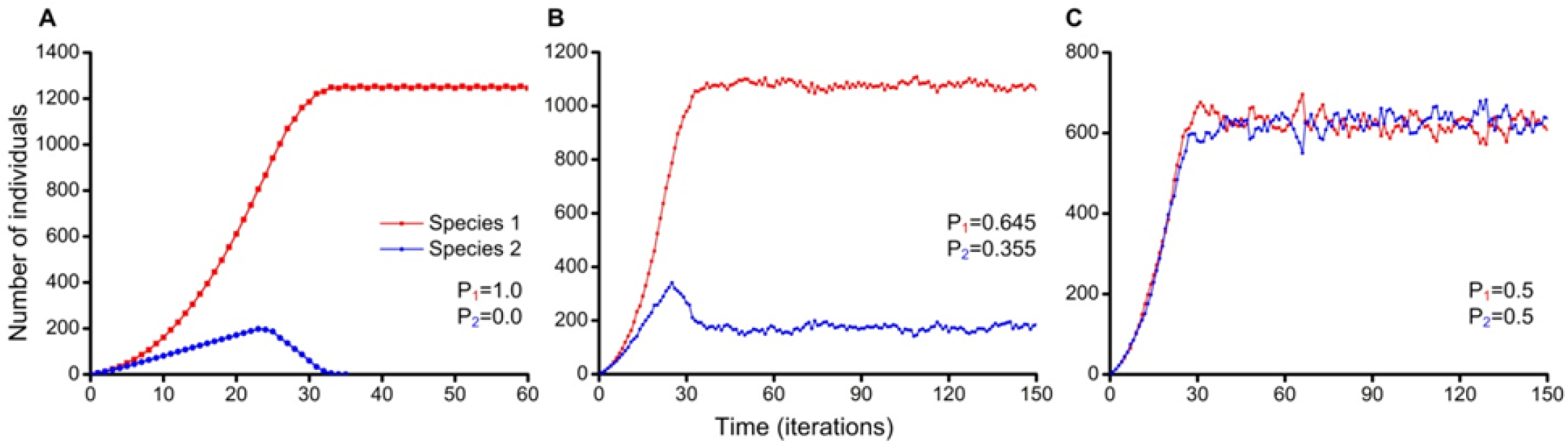
Population dynamics of two competing species. Colonization of free habitat which consists of 50x50 microhabitats started from a single individual of Species 1 and a single individual of Species 2. Parameter P_1_ reflects fitness of Species 1 and parameter P_2_ reflects fitness of Species 2. (A) The completely deterministic case of competitive exclusion when Species 1 excludes Species 2 (Movie S1). Species difference in fitness equals 100%. (B) Species 1 and Species 2 stably coexist despite the difference in fitness which equals 29% (Movie S2). (C) Species 1 and Species 2 are identical in all parameters and they stably coexist with close numbers of individuals (Movie S3). Species difference in fitness equals 0%.

The case of competitive exclusion closely reproduces dynamics from the Gause’s experiments with Paramecium aurelia and Paramecium caudatum, when they were cultivated in the mixed population (Fig. 4A and Movie S1) (Gause, 1934). In this case the complete competitors cannot coexist and only the fittest Species 1 survives. Collision of population waves of the two species in the habitat leads to annihilation of population waves of the Species 2. In Fig. 4B and Movie S2 we show how two completely competing species coexist despite 29% difference in fitness. Colliding population waves demonstrate diffusion-like behaviour. In the case when competing species are completely identical they coexist with close numbers of individuals (Fig. 4C and Movie S3). Colliding population waves also demonstrate diffusion-like behaviour.

Figure 4B and Movie S2 demonstrate a possibility of coexistence of competing species with 29% difference in fitness. A procedure of determination of a winner in each conflict of interest between individuals of competing species has a probabilistic nature (Fig. 1). In order to check the influence of the lattice size in the contribution of stochastic fluctuations on results of competition, we investigated competition of two species with *the same fitness* in four small ecosystems with different lattice sizes (Fig. 5).

**Figure 5.**
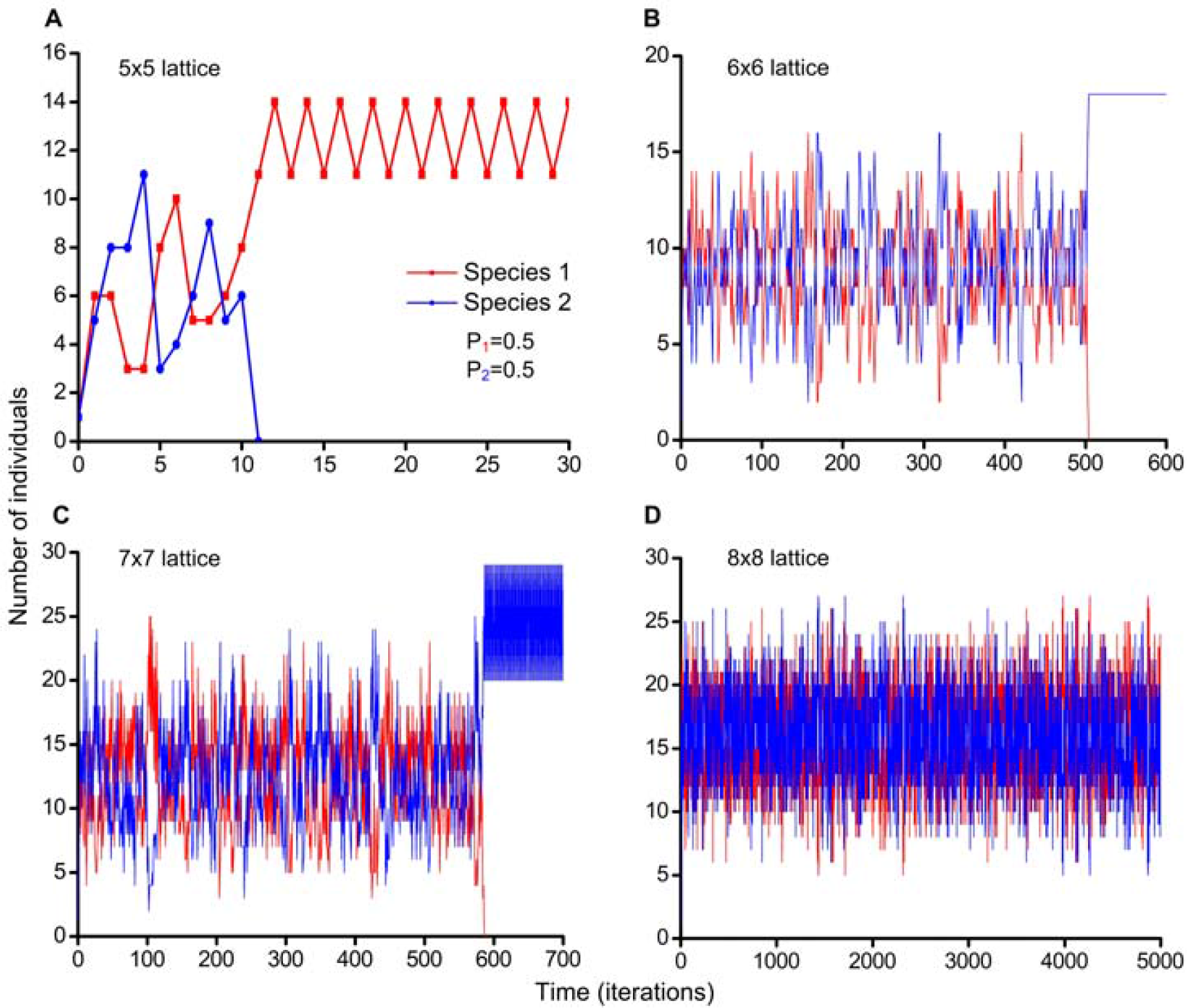
Influence of the habitat size in contribution of stochastic fluctuations on coexistence of two identical resource competitors. Colonization of free habitat started from single individuals of each species. Parameter P_1_ = 0.5 reflects fitness of the Species 1 and parameter P_2_ = 0.5 reflects fitness of Species 2. Raw data is available on figshare in Table S1 (https://doi.org/10.6084/m9.figshare.4903013.v1). (A) The Species 1 excludes the Species 2. The lattice consists of 5x5 sites. (B) The Species 2 excludes the Species 1. The lattice consists of 6x6 sites. (C) The Species 2 excludes the Species 1. The lattice consists of 7x7 sites. (D) The Species 1 and the Species 2 stably coexist. The lattice consists of 8x8 sites.

On the 5x5 lattice competitive exclusion of one of the species was observed already at 11-th iteration (Fig. 5A). On the 6x6 lattice competitive exclusion of one of the species took 504 iterations (Fig. 5B). On the 7x7 lattice the competitive exclusion required 586 iterations (Fig. 5C). On the 8x8 lattice the competitive exclusion was not detected during 5000 iterations (Fig. 5D). The obtained results demonstrate that the smaller the size of the lattice, the more stochastic fluctuations of competition outcomes appear in favor of one of the species. This individual-based model reproduces a phenomenon of the neutral genetic drift (Turner and Miller, 2015), based on stochastic fluctuations similar to the Moran Process (Moran, 1958). In small ecosystems, even in the absence of a selection, stochastic fluctuations may drive one of the species to extinction. Let us give a simple analogy with symmetrical coin which if is tossed only several times can show an occasional repetition of the same side. However, if this procedure will be repeated one million times, than the result will differ slightly from 50% for the both sides of the coin. Increasing of the lattice size leads to increasing in the number of cases of local competition, and, as a consequence, the contribution of probabilistic fluctuations in favor of one of the species is leveled, and the revealed phenomena become more natural and lawful. Since with the increasing number of iterations and lattice sizes the revealed results become more objective, more consistent with laws of nature, we repeated the studies presented in Fig. 4 with increased number of iterations from 150 up to 10000 and the lattice size from 50x50 to 300x300 sites (Figs. 6-9). These refinement experiments were carried out as Monte Carlo simulations with 200 trials in each case. In order to populate the lattice on the initial iteration, it was iteratively populated with probability 0.5% for each site before the criterion is reached – 625 individuals on the lattice 50x50 sites (Figs. 3B and 8) and 22500 individuals on the lattice 300x300 sites (Fig. 9) of each of two competing species.

**Figure 6.**
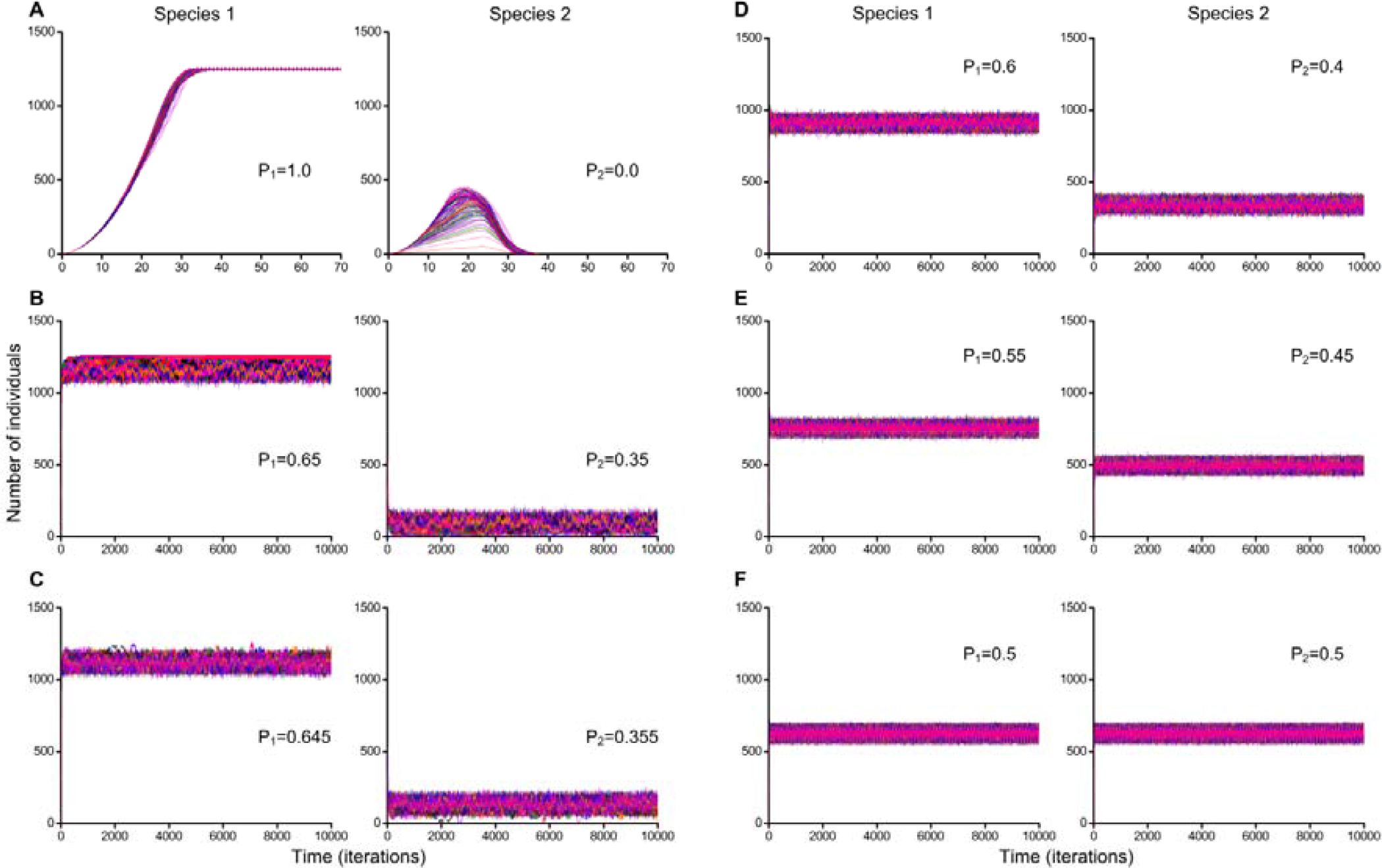
Monte Carlo simulations of two species competition with colonization of free habitat consisted of 50x50 sites. Colonization of free habitat started from single individuals of each species. The number of Monte Carlo simulations equals 200. Parameter P_1_ reflects fitness of Species 1 and parameter P_2_ reflects fitness of Species 2. Raw data is available on figshare in Table S2 (https://doi.org/10.6084/m9.figshare.4902986). (A) Competitive exclusion of the recessive Species 2 was observed in all cases. Species difference in fitness equals 100%. (B) There were 78 cases of competitive exclusion and 122 cases of competitive coexistence. Species difference in fitness equals 30%. (C-F) Competing species stably coexisted in all cases. Species differences in fitness equal 29% (C), 20% (D), 10% (E) and 0% (F).

**Figure 7.**
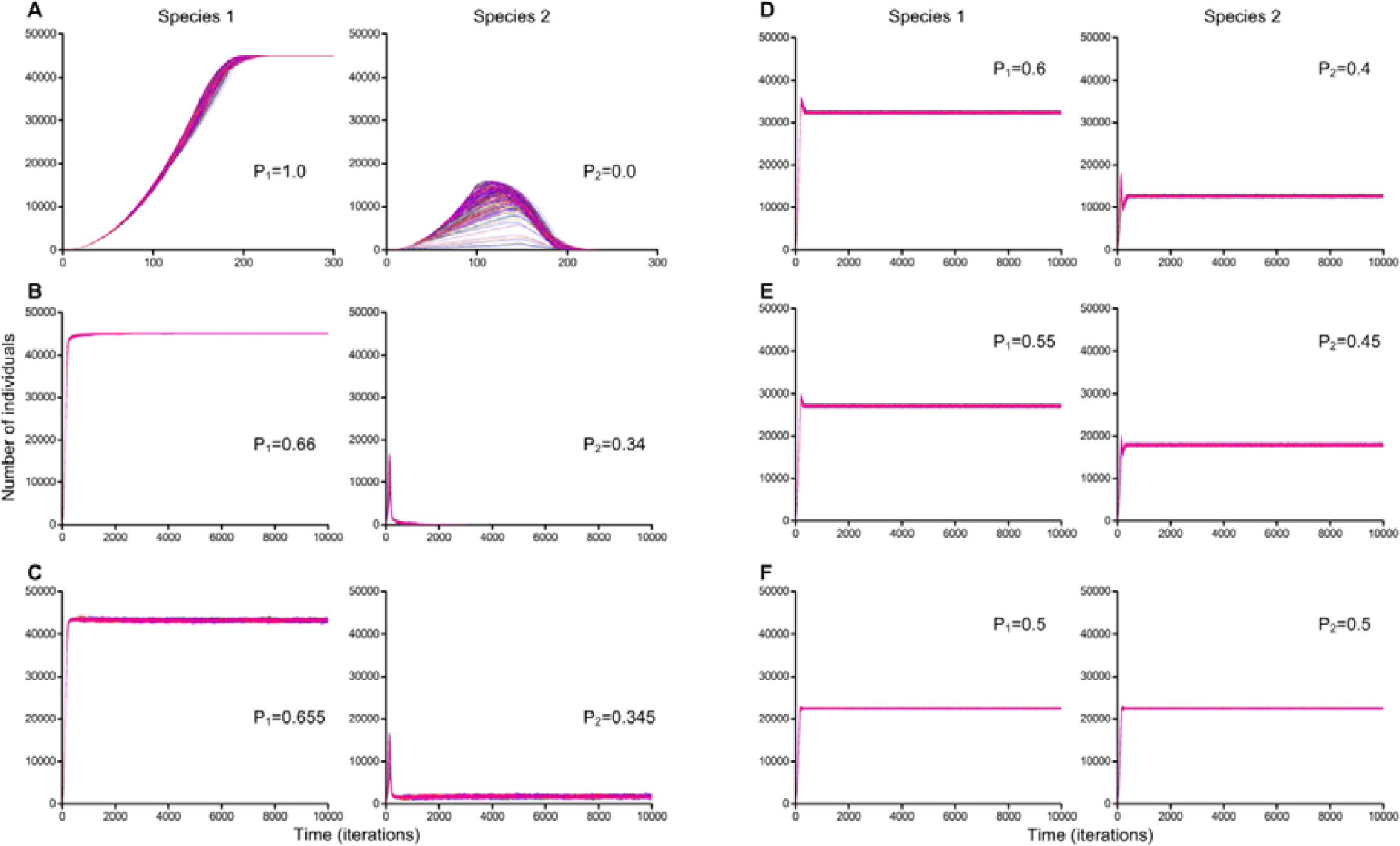
Monte Carlo simulations of two species competition with colonization of free habitat consisted of 300x300 sites. Colonization of free habitat started from single individuals of each species. The number of Monte Carlo simulations equals 200. Parameter P_1_ reflects fitness of the Species 1 and parameter P_2_ reflects fitness of the Species 2. Raw data is available on figshare in Table S3 (https://doi.org/10.6084/m9.figshare.4903016.v1). (A, B) Competitive exclusion of the recessive Species 2 was observed in all cases. Species differences in fitness equal 100% (A) and 32% (B). (C-F) Stable coexistence of competing species was observed in all cases. Species differences in fitness equal 31% (C), 20% (D), 10% (E) and 0% (F).

**Figure 8.**
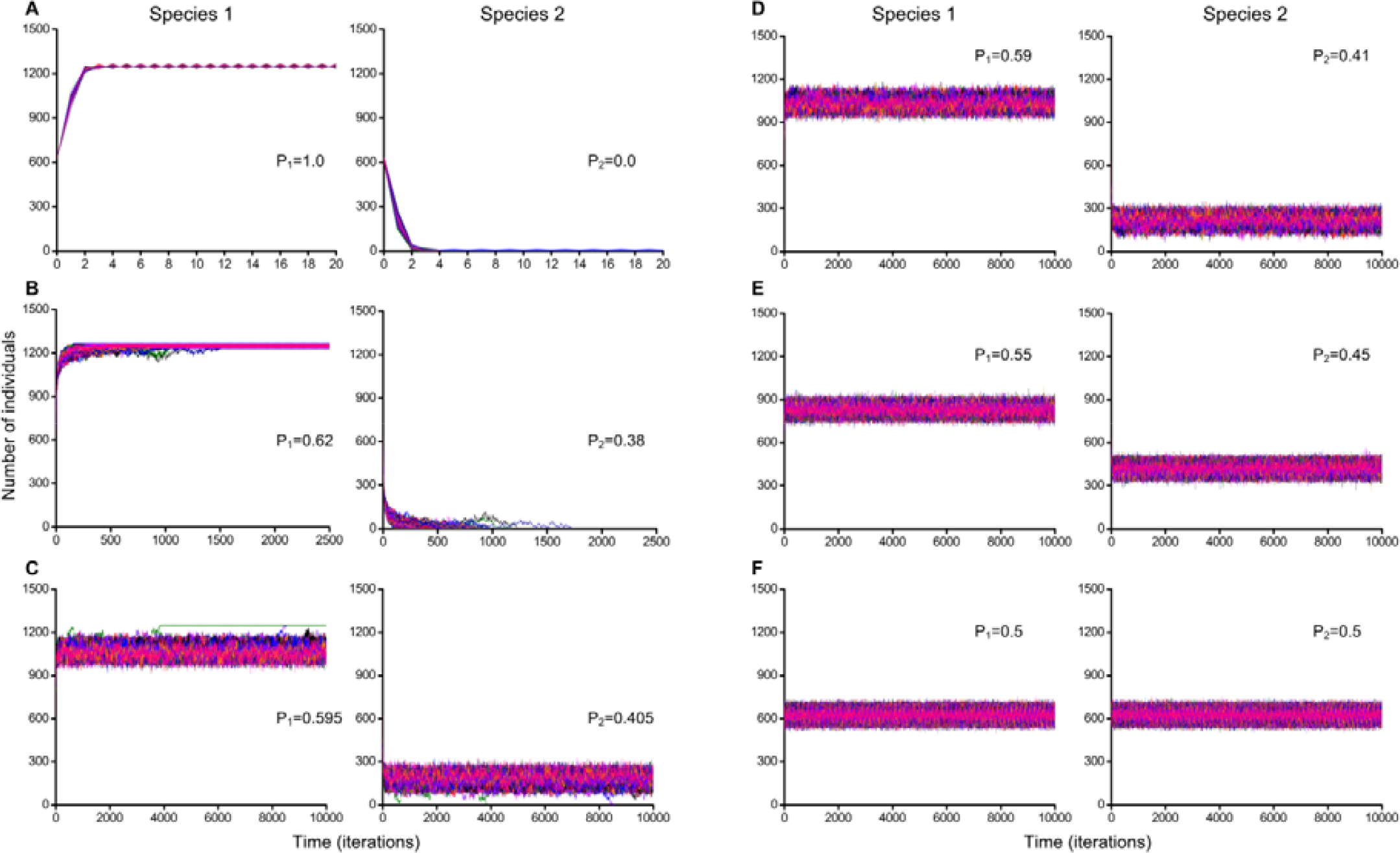
Monte Carlo simulations of two species competition when the habitat is probabilistically populated by individuals on initial iteration. Cellular automata field was probabilistically populated by 625 individuals of each of two competing species on initial iteration. The habitat consists of 50x50 microhabitats where 25% of the territory is occupied by individuals of the Species 1, 25% of the territory is occupied by individuals of the Species 2 and 50% of the territory is free. The number of Monte Carlo simulations equals 200. Parameter P_1_ reflects fitness of the Species 1 and parameter P_2_ reflects fitness of the Species 2. Raw data is available on figshare in Table S4 (https://doi.org/10.6084/m9.figshare.4903028.v1). (A) There were 198 cases of competitive exclusion and 2 cases of competitive coexistence. Species difference in fitness equals 100%. (B) Competitive exclusion was observed in all cases. Species difference in fitness equals 24%. (C) There were 2 cases of competitive exclusion and 198 cases of competitive coexistence. Species difference in fitness equals 19%. (D-F) Stable coexistence of competing species was observed in all cases. Species differences in fitness equal 18% (D), 10% (E) and 0% (F).

**Figure 9.**
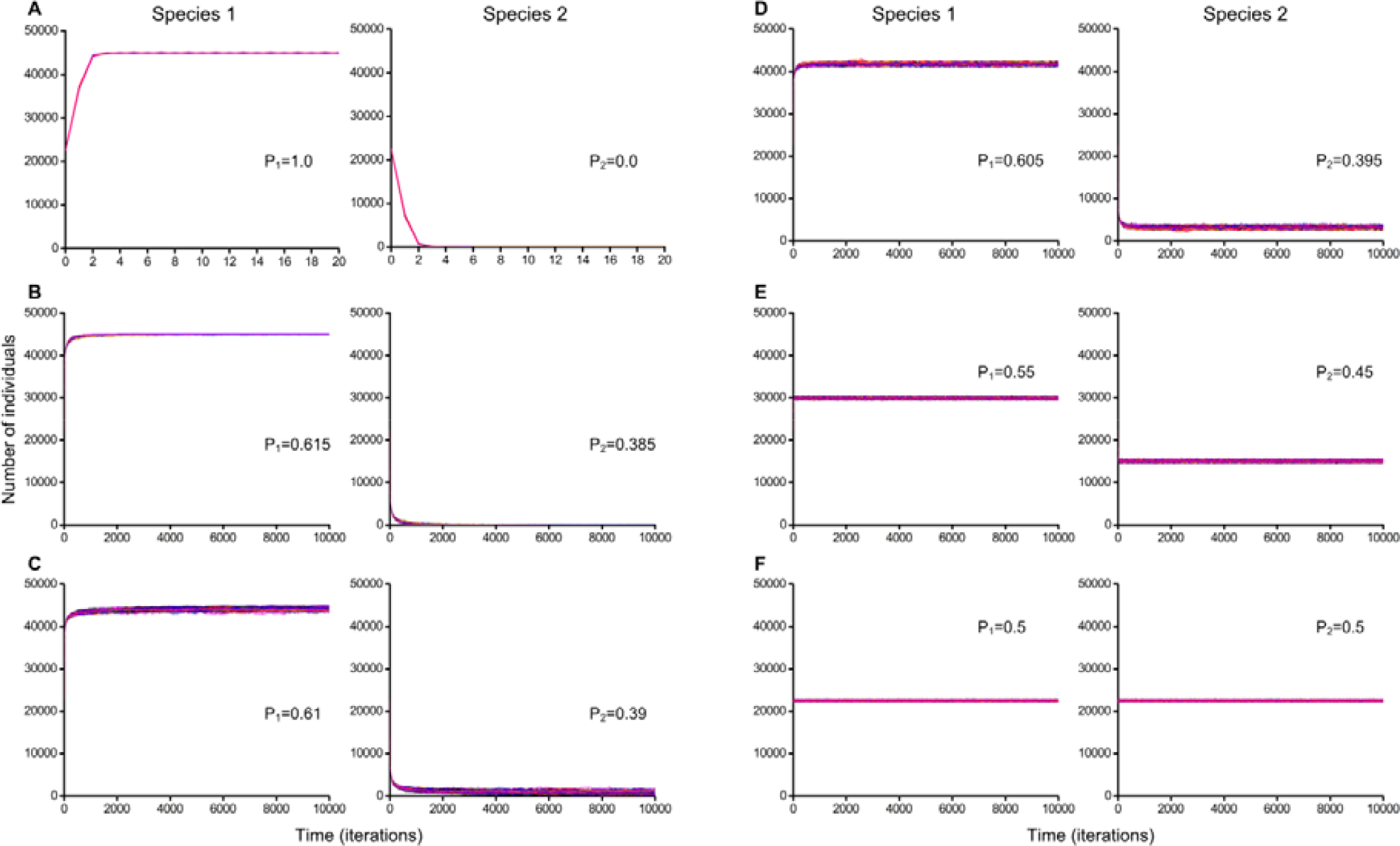
Monte Carlo simulations of two species competition when the habitat is probabilistically populated by individuals on initial iteration. Cellular automata field was probabilistically populated by 22500 individuals of each of two competing species on initial iteration. The habitat consists of 300x300 microhabitats where 25% of the territory is occupied by individuals of the Species 1, 25% of the territory is occupied by individuals of the Species 2 and 50% of the territory is free. The number of Monte Carlo simulations equals 200. Parameter P_1_ reflects fitness of the Species 1 and parameter P_2_ reflects fitness of the Species 2. Raw data is available on figshare in Table S5 (https://doi.org/10.6084/m9.figshare.4903031.v1). (A) There were 197 cases of competitive exclusion and 3 cases of competitive coexistence. Species difference in fitness equals 100%. (B) There were 195 cases of competitive exclusion and 5 cases of competitive coexistence. Species difference in fitness equals 23%. (C-F) Stable coexistence of competing species was observed in all cases. Species differences in fitness equal 22% (C), 21% (D), 10% (E) and 0% (F).

Using the Monte Carlo method, we have investigated the influence of different initial positioning of individuals on the lattice (Figs. 6-9). In Figs. 6 and 7 every Monte Carlo simulation consisted of 200 repeated experiments with different initial positioning of a single individual of the Species 1 and a single individual of the Species 2 on the 50x50 lattice (Fig. 6) and on the 300x300 lattice (Fig. 7), respectively.

In the case, when competing species had a maximum difference in fitness (P_1_ = 1.0; P_2_ = 0.0), then the Species 1 always outcompeted the Species 2 (Fig. 6A). This case does not contradict with the listed definitions of the competitive exclusion principle. In the case, when P_1_ = 0.65 and P_2_ = 0.35 there were 78 cases of competitive exclusion and 122 cases of competitive coexistence (Fig. 6B). When P_1_ = 0.645 and P_2_ = 0.355, i.e. the difference in fitness is 29%, competing species stably coexisted in all cases despite of different initial positioning of individuals of competing species in the habitat (Fig. 6C). At smaller differences in fitness the competing species also coexisted and their difference in number of individuals was smaller (Fig. 6C-F). These cases (Fig. 6C-F) demonstrate the validity of our hypothesis that two completely competing species can coexist with smaller difference in fitness despite of different initial positioning of individuals of competing species in the habitat.

Further, we conducted similar experiments for the larger lattice consisting of 300x300 sites (Fig. 7). We showed that due to increasing of the lattice size, competing species could coexist regardless initial placement of individuals with larger fitness difference of 31% (Fig. 7C). On the lattice of 50x50 sites it was 29%. Thus the influence of stochastic fluctuations in result of competitive interactions has been decreased noticeably. In the case demonstrated on Fig. 7F, the species have identical characteristics and have little difference in number after occupation of the habitat.

In Fig. 8 every Monte Carlo simulation consists of 200 repeated experiments with probabilistic positioning of 625 individuals of each species on the 50x50 lattice. An example of the initial pattern is shown in Fig. 3B. We discovered realization of a case of coexistence of complete competitors in 2 cases from 200 Monte Carlo experiments (Fig. 8A, Movie S4, Raw data in Table S4 - https://doi.org/10.6084/m9.figshare.4903028.v1). As a result of random initial location of individuals on the lattice, a particular pattern was realized that allowed the weak species to avoid a direct competition for resources with individuals of the dominant species. However, in such rare situation, the number of individuals of the recessive Species 2 was very small and fluctuated periodically between 1 and 6 – 1, 6, 1, 6, etc. This recessive species was on the verge of extinction, although it did not disappear. It is interesting to see how one recessive individual may successfully oppose 1246 individuals of the dominant species.

Increasing the habitat size from 50x50 to 300x300 resulted in an increase in the number of coexistence cases, – species always coexisted with differences in fitness equal to 29% (Fig. 6C) and 31% (Fig. 7C), respectively.

Next, we compared the cases in Fig. 6C with 2 individuals at the initial iteration and the cases in Fig. 8D with 625 individuals of each competing species. The habitat size had the same size – 50x50. In Fig. 6C the maximum value of the parameter P at which the species stably coexist equals 0.29 (29% difference in fitness) and in Fig. 8D the parameter P equals 0.18 (18% difference in fitness). Thus, when on initial iteration the habitat was populated only by 2 individuals there were the larger number of cases of competitive coexistence than when 50% of the habitat was initially populated (Figs. 3, 6, 8). At the identical sizes of the habitat, the species coexisted better when their competition began with two individuals at the initial iteration.

In Fig. 9 every Monte Carlo simulation consists of 200 repeated experiments with probabilistic positioning of 22500 individuals of the Species 1 and 22500 individuals of the Species 2 on the 300x300 lattice.

In the case shown in Fig. 9A there were 3 cases of competitive coexistence as a result of implementation of the mechanism of avoiding a direct conflict of interest, which was described for the case in Fig. 8A and Movie S4.

Increasing of the habitat size has led to increasing the number of cases of coexistence of competing species in conditions of direct competition for resources (Figs. 8D, 9C). In Fig. 8D species coexisted in all 200 cases at P = 0.18 (18% difference in fitness), and in Fig. 9C at P = 0.22 (22% difference in fitness).

Figs. 8C and 9B demonstrate cases of competitive exclusion when species had minimal differences in fitness. Decreasing the difference in fitness at 1% led to stable coexistence in all cases (Figs. 8D and 9C).

Sustainable coexistence of complete competitors with fitness inequality found here is an additional argument in favor that the classic formulations of the competitive exclusion principle should be reformulated in the direction of a greater number of conditions that must be met for implementation of the principle.

## CONCLUSIONS

This study was carried out using our extended individual-based cellular automata model of resource competition between two species. Previously we proved that complete competitors may coexist on one limiting resource regardless their 100% difference in fitness (Kalmykov and Kalmykov, 2013, 2015a). However, these results depended on initial positioning of individuals of the competing species in the habitat. Here we demonstrated that reducing the difference in fitness may help to overcome this dependence. We have found theoretically a new fact that two aggressively propagating competitors, which are identical consumers, can stably coexist on one limiting resource in one limited, stable and homogeneous habitat, when one species has some advantage in fitness over the other and all other characteristics are equal, in particular any trade-offs and cooperations are absent. This competitive coexistence is carried out regardless of initial location of individuals in the habitat, and, consequently, *the hypothesis of this study has been confirmed –* completely competing species may coexist with less than 100% difference in fitness regardless of different initial location of competing individuals in the ecosystem. The revealed coexistence of complete competitors additionally supports our reformulation of the competitive exclusion principle, which permited to resolve the biodiversity paradox.

## Acknowledgements

This research was supported by the Russian Foundation for Basic Research (No. 16-31-00516).

## REFERENCES

Bennett, A.E., Bever, J.D., 2009. Trade-offs between arbuscular mycorrhizal fungal competitive ability and host growth promotion in Plantago lanceolata. Oecologia 160, 807–816.

Cincotta, R.P., Wisnewski, J., Engelman, R., 2000. Human population in the biodiversity hotspots. Nature 404, 990–992.

Clark, J.S., 2009. Beyond neutral science. Trends in Ecology & Evolution 24, 8–15.

Clark, J.S., 2012. The coherence problem with the Unified Neutral Theory of Biodiversity. Trends Ecol Evol 27, 198–202.

Darlington, P.J., Jr., 1972. Competition, competitive repulsion, and coexistence. Proc Natl Acad Sci U S A 69, 3151–3155.

Dollhopf, S.L., Pariseau, M.L., Hashsham, S.A., Tiedje, J.M., 2003. Competitive and Cooperative Interactions Affecting a Fermentative Spirochete in Anaerobic Chemostats. Microbial Ecology 46, 1–11.

Ehrlich, P.R., Ehrlich, A.H., 2008. The Dominant Animal: Human Evolution and the Environment. Island Press.

Gause, G.F., 1934. The struggle for existence. Williams & Wilkins, Baltimore 163 pp.

Grubb, P.J., 1977. The maintenance of species-richness in plant communities: the importance of the regeneration niche. Biological Reviews 52, 107–145.

Hardin, G., 1959. Nature and man’s fate. Rinehart, New York.

Hardin, G., 1960. The Competitive Exclusion Principle. Science 131, 1292–1297.

Hardin, G., 1968. The Tragedy of the Commons. Science 162, 1243–1248.

Hastings, A., 1980. Disturbance, coexistence, history, and competition for space. Theoretical Population Biology 18, 363–373.

Hubbell, S.P., 2001. The unified neutral theory of biodiversity and biogeography. Princeton University Press, Princeton, N.J.; Oxford xiv, 375 p. pp.

Huston, M., DeAngelis, D., Post, W., 1988. New Computer Models Unify Ecological Theory: Computer simulations show that many ecological patterns can be explained by interactions among individual organisms. BioScience 38, 682–691.

Hutchinson, G.E., 1961. The paradox of the plankton. The American Naturalist 95, 137–145.

Kalmykov, L.V., Kalmykov, V.L., 2013. Verification and reformulation of the competitive exclusion principle. Chaos, Solitons & Fractals 56, 124–131.

Kalmykov, L.V., Kalmykov, V.L., 2015a. A Solution to the Biodiversity Paradox by Logical Deterministic Cellular Automata. Acta Biotheor 63, 203–221.

Kalmykov, L.V., Kalmykov, V.L., 2015b. A white-box model of S-shaped and double S-shaped single-species population growth. PeerJ 3, e948.

Kalmykov, V.L., Kalmykov, L.V., 2016. On ecological modelling problems in the context of resolving the biodiversity paradox. Ecological Modelling 329, 1–4.

Krinsky, V.I., 1984. Autowaves: Results, Problems, Outlooks, in: Krinsky, V.I. (ed.), Self-Organization Autowaves and Structures Far from Equilibrium: Proceedings of an International Sympusium Pushchino, USSR, July 18–23, 1983. Springer Berlin Heidelberg, Berlin, Heidelberg, pp. 9–19.

Lehman, C.L., Tilman, D., 1997. Spatial ecology: the role of space in population dynamics and interspecific interactions, in: Tilman, D., Kareiva, P.M. (eds.), Monographs in Population Biology Princeton University Press, Vol. 30. Princeton University Press Princeton, NJ, pp. 185–203.

Levin, S.A., 1970. Community Equilibria and Stability, and an Extension of the Competitive Exclusion Principle. The American Naturalist 104, 413–423.

MacArthur, R., Levins, R., 1964. Competition, Habitat Selection, and Character Displacement in a Patchy Environment. Proceedings of the National Academy of Sciences of the United States of America 51, 1207–1210.

Moran, P.A.P., 1958. Random processes in genetics. Mathematical Proceedings of the Cambridge Philosophical Society 54, 60–71.

Narwani, A., Berthin, J., Mazumder, A., 2009. Relative importance of endogenous and exogenous mechanisms in maintaining phytoplankton species diversity. Ecoscience 16, 429–440.

Nowak, M.A., 2006. Five Rules for the Evolution of Cooperation. Science 314, 1560–1563.

Palmer, M.W., 1994. Variation in Species Richness - Towards a Unification of Hypotheses. Folia Geobot Phytotx 29, 511–530.

Reichenbach, T., Mobilia, M., Frey, E., 2007. Mobility promotes and jeopardizes biodiversity in rock-paper-scissors games. Nature 448, 1046–1049.

Rosindell, J., Hubbell, S.P., He, F., Harmon, L.J., Etienne, R.S., The case for ecological neutral theory. Trends in Ecology & Evolution 27, 203–208.

Silvertown, J., Holtier, S., Johnson, J., Dale, P., 1992. Cellular Automaton Models of Interspecific Competition for Space - the Effect of Pattern on Process. Journal of Ecology 80, 527–534.

Sommer, U., 1999. Ecology - Competition and coexistence. Nature 402, 366–367.

Spencer, H., 1864. The Principles of Biology. Williams and Norgate, London.

Tansley, A.G., 1935. The Use and Abuse of Vegetational Concepts and Terms. Ecology 16, 284–307.

Tilman, D., 1987. The importance of the mechanisms of interspecific competition. The American Naturalist 129, 769–774.

Turner, A.J., Miller, J.F., 2015. Neutral genetic drift: an investigation using Cartesian Genetic Programming. Genetic Programming and Evolvable Machines 16, 531–558.

Vitousek, P.M., Mooney, H.A., Lubchenco, J., Melillo, J.M., 1997. Human Domination of Earth’s Ecosystems. Science 277, 494–499.

Watt, A.S., 1947. Pattern and Process in the Plant Community. Journal of Ecology 35, 1–22.

Wellborn, G.A., 2002. Trade-Off between Competitive Ability and Antipredator Adaptation in a Freshwater Amphipod Species Complex. Ecology 83, 129–136.

Whitfield, J., 2002. Neutrality versus the niche. Nature 417, 480–481.

Wiener, N., Rosenblueth, A., 1946. The mathematical formulation of the problem of conduction of impulses in a network of connected excitable elements, specifically in cardiac muscle. Archivos del Instituto de Cardiologia de Mexico 16, 205–265.

Zaikin, A.N., Zhabotinsky, A.M., 1970. Concentration Wave Propagation in Two-dimensional Liquid-phase Self-oscillating System. Nature 225, 535–537.

